# Charge-based fingerprinting of unlabeled full-length proteins using an aerolysin nanopore

**DOI:** 10.1101/2025.01.13.632743

**Authors:** Verena Rukes, Evita Norkute, Georges Barnikol, Jingze Duan, Jiajie Gao, Chan Cao

## Abstract

Proteins play essential roles in cellular processes and are involved in numerous diseases, driving the need for efficient proteoform identification. While nanopore technology was highly successful for DNA sequencing, this approach has not yet delivered its full potential in protein identification. Here, we demonstrate the capabilities of an aerolysin nanopore for direct identification of a range of unlabeled full-length proteins. Combining low pH and guanidinium chloride, we generate a strong electroosmotic flow that enables an efficient capture and translocation, resulting in distinct fingerprints from single-protein translocations. Using machine learning classifier, we achieve 80% accuracy in distinguishing seven related proteins with 38%–70% pairwise sequence identities. Differences in fingerprints largely reflect the distribution of positive charges in the protein sequences, providing a rational basis for the observed sensitivity. With further development, fingerprint-predictions could allow to infer *de novo* protein sequence information from single-molecule data, offering a powerful tool for proteomics.

## Introduction

Proteins are the most abundant macromolecules in living systems and fulfill a plethora of functions such as providing cell structure, and signaling pathways, or driving metabolic processes^1,2^. They are highly diverse, accounting for millions of human proteoforms. Many of these are the product of post transcriptional alterations in the cell that dictate protein function and cannot be accessed through the ∼20,000 protein-encoding genes^3–5^. This is why tools that directly access the protein level have great significance e.g. in diagnostics. Currently, protein analysis is mostly based on mass spectrometry (MS), which despite much optimization^6^ is still accompanied by drawbacks such as high costs, limited dynamic range, and detection limit as well as a limited reading length^7–9^. This has resulted in a rising interest in the development of single-molecule techniques to expand the repertoire of protein-analysis tools.

Recent advancements have highlighted nanopores as promising tools for protein analysis, using voltage-driven ionic currents through a nanometer-scale pore^10,11^. As analytes interact with the pore, the ionic currents are modulated^12^, which is commercialized for DNA sequencing^13^. Notably, nanopores can measure single-molecule interactions and are compatible with mixture samples. To tailor the method to protein analysis, several approaches are being demonstrated, such as using DNA^14–16^ or protein^17,18^ motor enzymes to slowly thread peptides/proteins though nanopores. Another strategy is to fragment proteins and fingerprint the peptide spectra similar to bottom-up MS^19–21^. However, these require sample processing steps like chemical linkage or protein digest, that will reduce yield and introduce uncertainty to the analysis.

A direct label-free approach can provide faster detection, specifically of low abundant species, which is desired in protein analysis. Conceptually, this can be achieved by directly threading the full-length proteins through the pore and identifying each protein based on prior knowledge of its current readout^22^. Important groundwork towards this goal has demonstrated that chemical denaturants can be used during nanopore measurements to weaken an analytes’ structure^23^, which is necessary to access sequence specific information. Since proteins are heterogeneously charged, it is not possible to rely solely on the electrophoretic force (EPF) as a driving force. This has been overcome by taking advantage of the Electroosmotic Flow (EOF), which can attract and transport analytes through nanopores independently of their charge. The EOF relies on the unequal transport of ions species of biological pores (illustrated in Fig. 1A), which is referred to as ion selectivity and is caused by the charge of the pore lumen. Using the EOF, proteins and peptides have been successfully translocated through nanopores^24–26^. Selectivity can be induced and tuned through mutations^25^, pH^27^, or the electrolyte composition. For instance, guanidinium chloride (GdmCl) has been shown to universally induce EOF in nanopores^28^.

**Figure 1.**
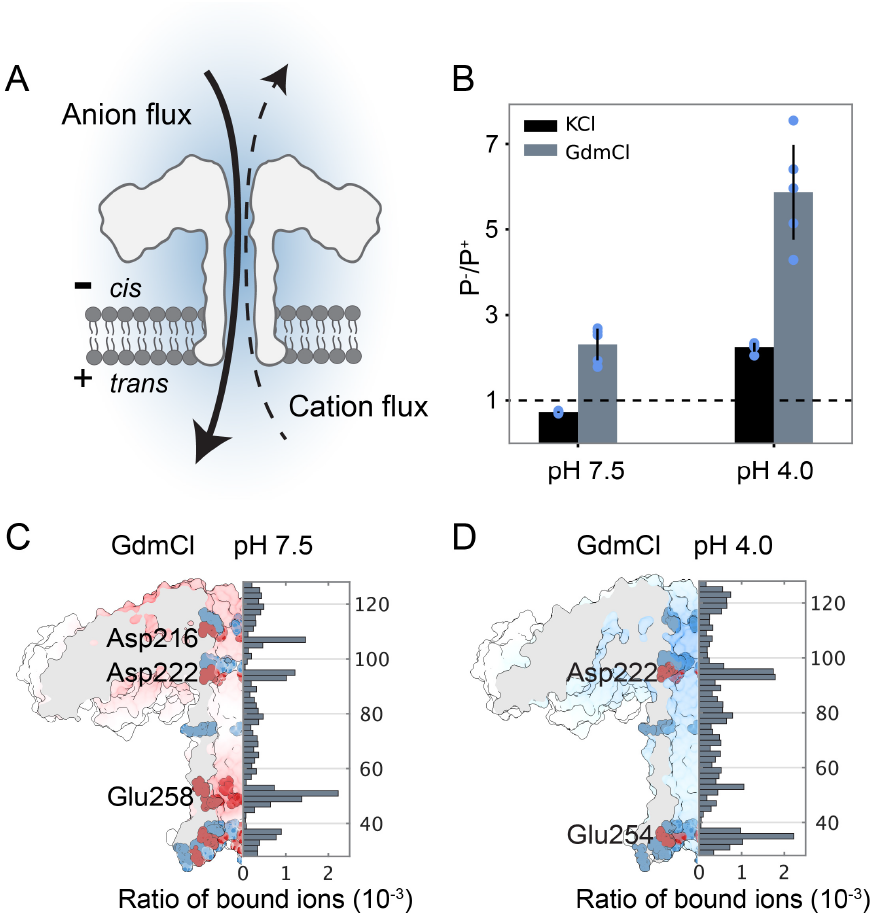
Enhancing ion selectivity by combining pH and GdmCl. (A) Illustration of anion selectivity in aerolysin. (B) Permeability ratios based on reversal potential measurements with aerolysin K238A in KCl or GdmCl at pH 7.5 and pH 4.0. Equal permeability is indicated with a dashed line (P^-^/P^+^ = 1). In each condition P^-^/P^+^ was averaged over > 4 independent measurements and standard deviation*s* are indicated. MD results for GdmCl at pH 7.5 (C) and pH 4.0 (D) are shown together with an illustration of the pore structure. Residues carrying negative and positive charge are marked in red and blue respectively. Bar plots illustrate the duration of Gdm^+^ binding to the pore surface normalized by the simulation time.

Full-length protein identification was recently demonstrated using alpha hemolysin (α-HL)^22^ or homologous pores (CytK)^25^. However, aerolysin is known for its high resolution when distinguishing analytes in free translocation^29,30^. Specifically, the aerolysin mutant K238A has been used for identification of site specific PTMs in peptides and direct decoding of data-storage polymers^31,32^. Due to its narrow cylindrical shape, aerolysin has been described as an ideal candidate for linear protein transport^33^. Additionally, single-file translocations of an unfolded protein has been reported with aerolysin^34^. However, to our knowledge, there has been no experimental and analytical investigation focused on establishing this nanopore for identification of full-length proteins.

In this work, we demonstrate the optimization of EOF and unfolding conditions in aerolysin for efficient translocation of full-length proteins. In an optimized condition of 3 M GdmCl at pH 4.0, we measured 7 proteins from the same Turandot family of *Drosophila Melanogaster* to explore the sensitivity of the system in an extreme case. At an optimal potential of 170 mV, machine learning classifiers were able to discriminate between these 7 analytes with 80% accuracy. We further investigated the dependence of the current output on the sequence of the analyte. A simplified biophysical model along with the experiments of control analytes demonstrated that patterns in the current readout can be attributed to the distribution of positive charges along the protein analyte sequences. Current-sequence dependency has not previously been explored for freely translocating unfolded proteins. However, establishing such rational connection is the first step to develop an accurate model, which could significantly enhance this approach and potentially enable *de novo* protein identification.

## Results

### Enhancing EOF in the aerolysin system

The central importance of the EOF for protein capture in nanopore systems is established. We thus characterized the impact of changes in the measurement condition to optimize the EOF in aerolysin K238A. While the structural integrity of the aerolysin pore is maintained up to 3 M GdmCl^35^, it is known that the EOF in nanopores is affected by both Gdm^+ 22,28^ and changes in pH^27^. Based on reversal potential measurements we calculated the ratio of permeability (*P*^*-*^*/P*^*+*^) between anions and cations through the pore (Fig. 1B and SI Fig. S1). This ratio is commonly used to determine the direction and relative magnitude of ion selectivity in biological nanopore systems^25,28,36–38^. Aerolysin K238A is slightly cation selective at neutral pH in KCl (*P*^*-*^*/P*^*+*^ = 0.73 ± 0.02). By changing the pH to 4.0 or changing the electrolyte to GdmCl, we observed a strong anion selectivity, resulting in *P*^*-*^*/P*^*+*^ = 2.24 ± 0.10 and *P*^*-*^*/P*^*+*^ = 2.31 ± 0.37, respectively. When combining both conditions (GdmCl at pH 4.0), an even larger selectivity is reached with *P*^*-*^*/P*^*+*^ = 5.87 ± 1.11 in aerolysin K238A. Thus, we establish the combination of low pH and GdmCl ions as a fast and simple strategy to turn a slightly cation selective pore system into a strongly anion selective one.

To investigate how or whether the binding of Gdm^+^ is impacted by the change in pH, we have performed all atoms molecular dynamics (MD) simulations with aerolysin K238A in KCl and GdmCl both at pH 7.5 and pH 4.0. Interaction spots of the ions with the pore were defined, by considering ions that enter the pore and monitoring the time that each ion type spends in close proximity along the surface of the pore lumen, similar as previously reported^28^. We then normalize by the duration of the trajectory (see details in Methods). Gdm^+^ showed specific interaction spots along the pore at both pH. At pH 7.5, Gdm^+^ lingered at residues Asp216, Asp222, and (most prominently) Glu258, resulting in distinct peaks seen in Fig. 1C. At pH 4.0, the peaks at residues Asp216 and Glu258 largely disappear. While Asp222 remained a site of interaction, the residence of Gdm^+^ at Glu254 increased (Fig. 1D). At either pH such peaks did not occur for Cl^-^ in the GdmCl condition or for the ions in a KCl condition (SI Fig. S2).

Overall, MD results – in agreement with experiments – showed that Gdm^+^ ions can bind the pore lumen both at pH 7.5, and at pH 4.0. At both pH, the Gdm^+^ interaction spots align well with negative charges in the lumen, showing the importance of electrostatic interactions for the binding. The combination of low pH and GdmCl results in more than double the *P*^*-*^*/P*^*+*^ without noticeable negative interference, therefore providing a simple strategy to produce extreme ion selectivity and EOF.

### Optimizing the measurement condition for protein translocation

We used Turandot A (TotA) from *Drosophila Melanogaster* to optimize the buffer condition for full length protein analysis with aerolysin K238A. At pH 7.5, TotA has a net charge of - 7.8 and is driven toward the pore by both the EPF and the EOF, while at pH 4.0 the protein’s net charge is + 8.0 and therefore the EOF is the only attracting force, opposing the EPF (Fig. 2A). The TotA protein has 115 amino acids (aa) and its native structure is composed of four α-helices^39^ (Fig. 2B). Notably, changing the pH from 7.5 to 4.0 does not impact on the Circular Dichroism (CD) spectrum of TotA, indicating no change in its structure (Fig. 2B). Additionally, CD showed that TotA is not fully unfolded at 2 M GdmCl and largely unfolded at 3 M GdmCl. Therefore, by measuring the protein at 2 M GdmCl and 3 M GdmCl both at pH 7.5 and pH 4.0, we investigate the importance of residual structure, and the EPF-EOF interplay in aerolysin measurements of a full-length protein. TotA interactions with the aerolysin K238A pore were measured under a voltage range from 100 mV to 210 mV applied to the *trans* compartment. In all conditions and at all voltages, the analyte was added to the *cis* compartment, which caused deep current blockages with less than 15% mean residual current and dwell times between 3 and 30 ms (Fig. 2C-E, SI Fig. S3).

**Figure 2.**
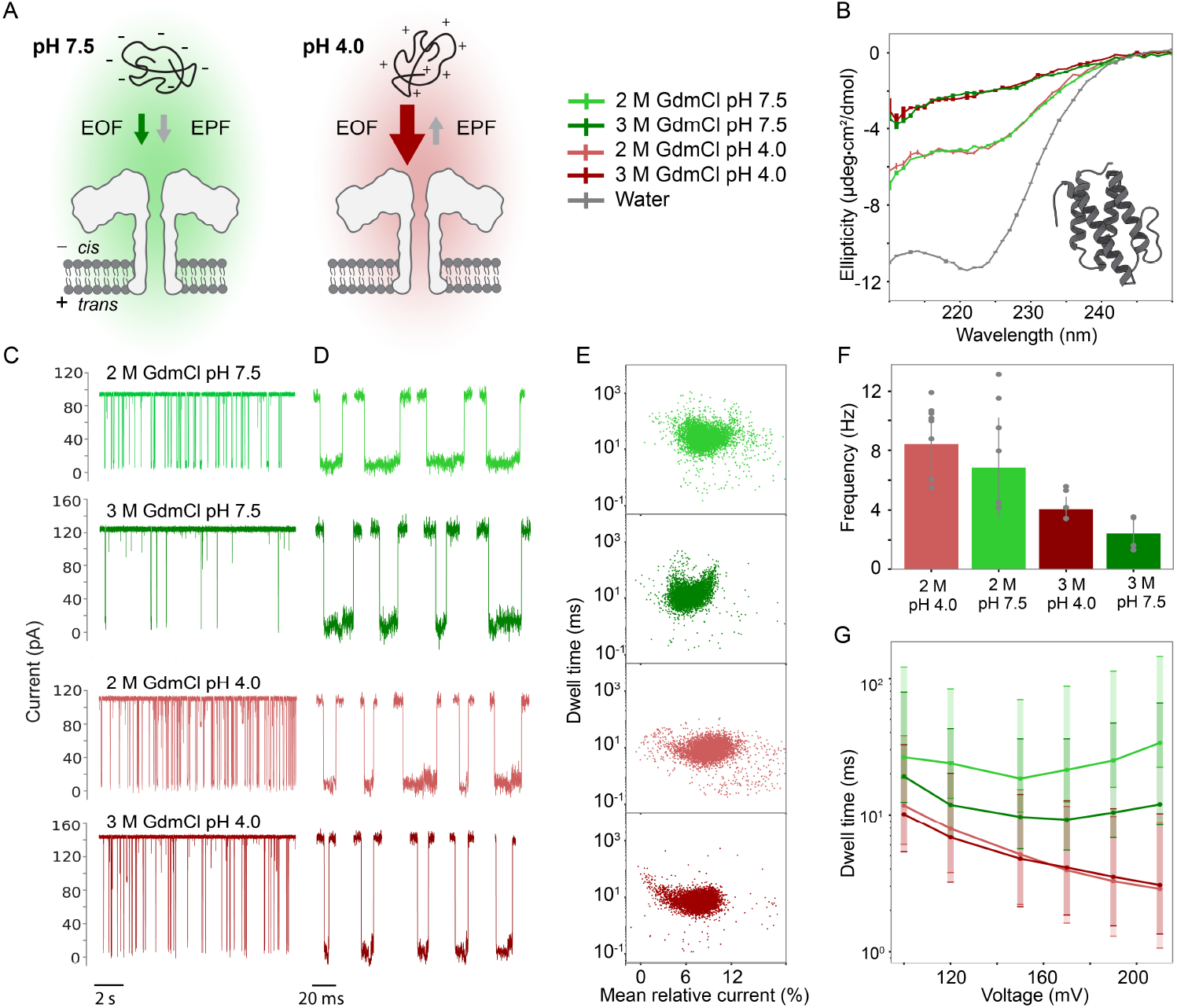
Optimizing conditions for protein translocations. (A) Illustration of the interplay of EOF and EPF in GdmCl at pH 7.5 (left) and pH 4.0 (right). (B) Results of CD measurements are averaged over 3 repeated measurements; the standard deviations are indicated. The TotA structure is shown (PDB: 8PBV). (C) 10 s of raw trace for each condition filtered to 1 kHz for illustration. (D) Example events filtered to 10 kHz for illustration. (E) Dwell time versus current scatterplots for individual events and each condition recorded at 120 mV. (F) Frequency of events calculated by fitting the interevent time of individual pores. Averaged over 8, 6, 5, and 4 recordings from left to right respectively weighted by the number of events in each recording, standard deviations are indicated. (G) The distribution of log dwell times for each condition at each voltage were fitted using a single exponentially modified gaussian distribution. The peak and the halfwidth of the fitted distributions are shown.

We found that generating a strong EOF even against the EPF (pH 4.0) is the more efficient strategy to capture and translocate the protein analyte, compared to a combination of EPF and lower EOF (pH 7.5). As shown in Fig. 2F, at 100 mV shifting the pH from 7.5 to 4.0 increased capture at both 2 M and 3 M GdmCl, confirming that there is a stronger driving force toward the pore at low pH, even against the EPF. More strikingly, at pH 7.5, the dwell times did not decrease monotonously with increasing voltages (Fig. 2G). We observed a turnover voltage of 150 mV in 2 M GdmCl, which shifted to 170 mV at 3 M GdmCl. Therefore, we cannot conclude that TotA can translocated the pore at pH 7.5. However, despite being captured against the EPF, the analyte did translocate the engineered aerolysin pore at pH 4.0 in both 2 M and 3 M GdmCl (Fig. 2G). Interestingly, the residual structure of TotA in 2 M GdmCl did not hinder its translocation. At pH 4.0, the current pattern of the events as well as the shape of the event populations was very similar in 2 M and 3 M GdmCl, further indicating that the change in structure did not impact the process of protein translocation.

Increasing the GdmCl concentration from 2 M to 3 M increased the current at both pH (SI Fig. S4). Since the EOF is a result of net ion translocation, it is expected to be stronger at higher current (given equal selectivity) – and thus larger at 3 M than at 2 M GdmCl. However, while dwell times were faster at pH 7.5 with increased salt concentration, they remained similar at pH 4.0. We also observed that the capture efficiency decreased with increasing GdmCl concentration. A likely cause is the more unfolded structure in 3 M GdmCl resulting in a larger radius of gyration and thereby lower diffusion coefficient in bulk. Following this evaluation, we chose a condition of 3 M GdmCl at pH 4.0 for further studies.

### Measurements of a protein family

To further explore the interplay of full-length proteins with aerolysin in the optimized condition (in 3 M GdmCl, pH 4.0), we measured six more members of the Turandot family protein at 100 mV – 210 mV (Fig. 3A-B). These analytes have large structural similarities and pairwise sequence identities between 38% - 70% (Fig. 3A, SI Fig. S5-6). The analytes produced event populations with 0-10% mean residual current at 120 mV and clear deviations in dwell time between the different proteins (Fig. 3C, Example raw traces in SI Fig. S7-9). For all 7 proteins, the population dwell time decreased with increasing voltage, confirming their translocation (Fig. 3D).

**Figure 3.**
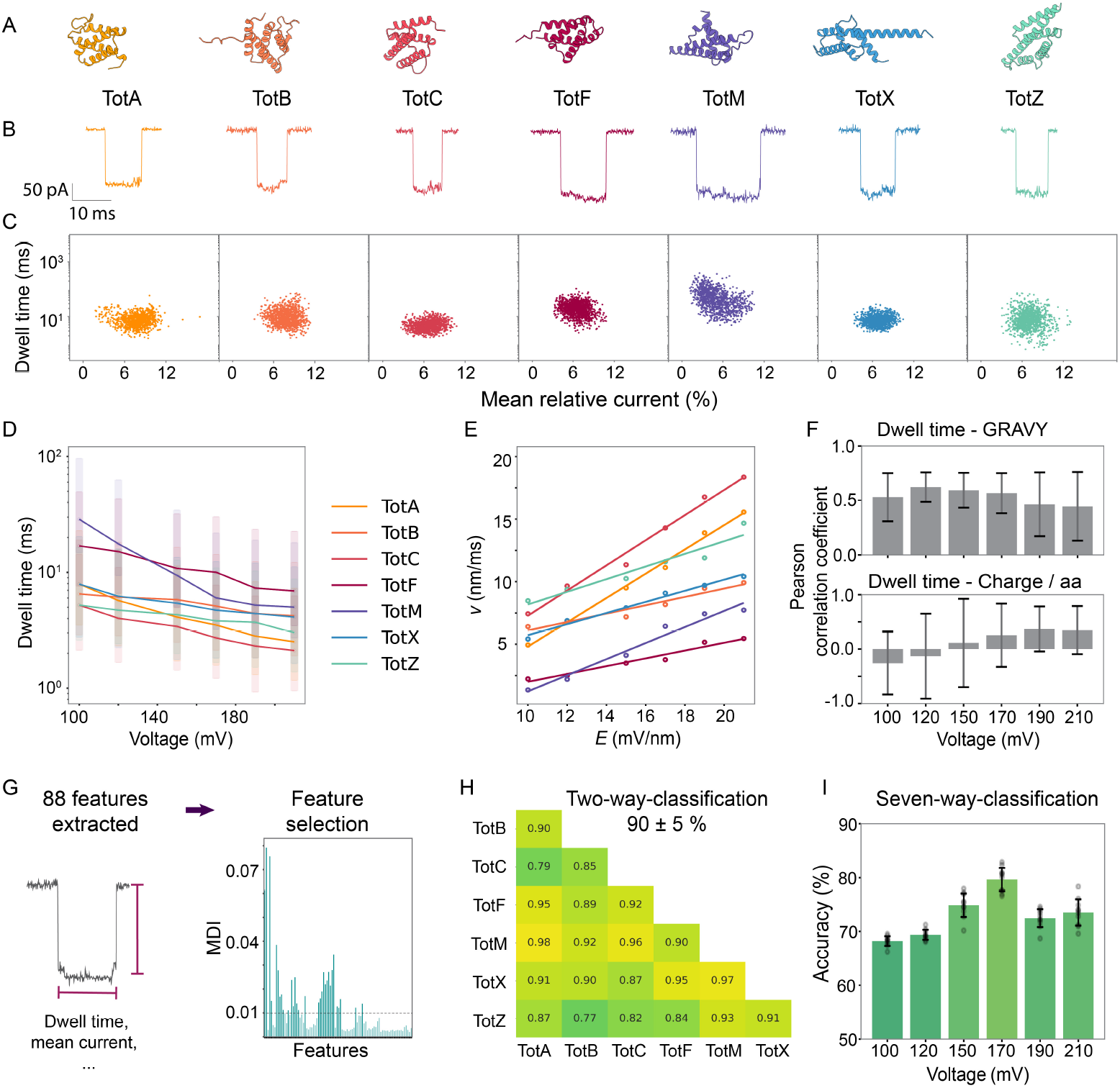
Measurements of related proteins. (A) Structural representation of each protein analyte (predicted using Alpha Fold 2^40^). (B) Example events collected at 120 mV for corresponding proteins. (C) Dwell time versus current scatterplots for individual events recorded at 120 mV of (left to right) TotA (yellow), TotB (orange), TotC (light red), TotF (dark red), TotM (purple), TotX (blue), TotZ (turquoise). (D) Population dwell times were fitted with the Fokker Planck equation for each protein and plotted against applied voltages. Maximum and halfwidth of the distributions are shown. (E) Velocities of each protein against electric field fitted with a linear regression. (F) Pearson correlations between the population dwell time and the GRAVY (top) and the charge normalized by the number of amino acids of each protein (bottom) cross different voltages. The p-values of the correlations are indicated as error bars. (G) Illustration of the feature extraction and reduction based on the mean decrease in impurity (MDI) for machine learning classification. (H) Two-way classification of the Tot family proteins using data recorded at 120 mV. (I) Accuracy of seven-way classifications of the Tot family proteins using data at different voltages. For each voltage 10 rounds of data selection and model training were performed and the average with indicated standard deviation is shown.

Translocation velocities (Fig. 3E, proteins length divided by the fitted dwell time) depended roughly linearly on the electric field, which is in alignment with expectations (SI Note). However, velocities are not consistent between analytes, since the dwell times poorly correlate with the analyte length (SI Fig. S10). This suggests that sequence-specific effects dominate the kinetics during protein translocation. The grand average hydropathy (GRAVY, calculated based on the Kyte-Doolittle hydropathy scale^41^) showed a weak positive correlation with the population dwell time across all voltages (Fig. 3F), while the charge per amino acid (aa) did not consistently correlate with the dwell times at all. Thus, while the velocity depended linearly on the electric field, the sequence of the analytes affected the absolute dwell times.

### Machine learning-based protein identification

Nanopore protein identification ultimately has the potential to identify even low abundant molecules in undefined samples. This requires the development of advanced analytical methods capable of identifying proteins through their single-molecule interactions. Machine learning classifiers have demonstrated strong performance in tackling this challenge^17,22,42^. We chose to describe each single-molecule interaction in 88 statistical features (Fig. 3G, SI Table 2). To allow maximal interpretability, we reduce the features by thresholding the mean decrease in impurity (MDI) in a random forest classification. This ensures that only features that are useful for identification were used further. From the MDI we observed that one of the most important features was the dwell time, showing that the protein-dependent variance in dwell time can ultimately be useful for identification. Additionally, the current and standard deviation in the first and last portions of the events were useful for identification (SI Fig. S11). We benchmark common classification algorithms and finally trained a voting classifier combining the top three models (see methods).

An initial two-way classification demonstrated clear differentiation between signals from the related proteins with an accuracy of 90 ± 5% at 120 mV (Fig. 3H). Expanding to a seven-way classification of all Turandot proteins yielded an accuracy of 69 ± 1%, reflecting the increased challenge of distinguishing a larger set of closely related analytes. Interestingly, we found that tuning the applied voltage affected classification accuracies (Fig. 3I). Increasing the voltage to 170 mV improved accuracies to 80 ± 2%, suggesting that the higher voltage amplified the variance between the protein-pore interactions, resulting in more pronounced differences in the single molecule fingerprints. However, further increase in voltage (>170 mV) reduced accuracies, likely due to the shorter event times at those voltages.

### Interpreting nanopore current based on protein sequences

To thoroughly analyze the current signature of each analyte, we randomly selected events and overlaid them in a single plot for detailed comparison (Fig. 4A, SI Fig. S12). We found that the current signatures of each analyte are directional which is most evident in the patterns for TotF, TotM, and TotX. This suggests that the analytes translocated the pore from the same terminus, although no charged tag was added to facilitate unidirectionality. For TotA, TotB and TotC, currents tended to slightly decrease around the middle of the event, while for TotX an increase in current appeared toward the middle of the interaction. TotF and TotM show a clear downward trend in current and TotZ had the most stable current. By comparing the protein sequences (SI Fig. S6, S13), we found that these distinct features of the current patterns could be explained by the distribution of positive charges along the proteins (Fig 4B), while volume exclusion (Fig 4C) was less relevant for the patterns.

**Figure 4.**
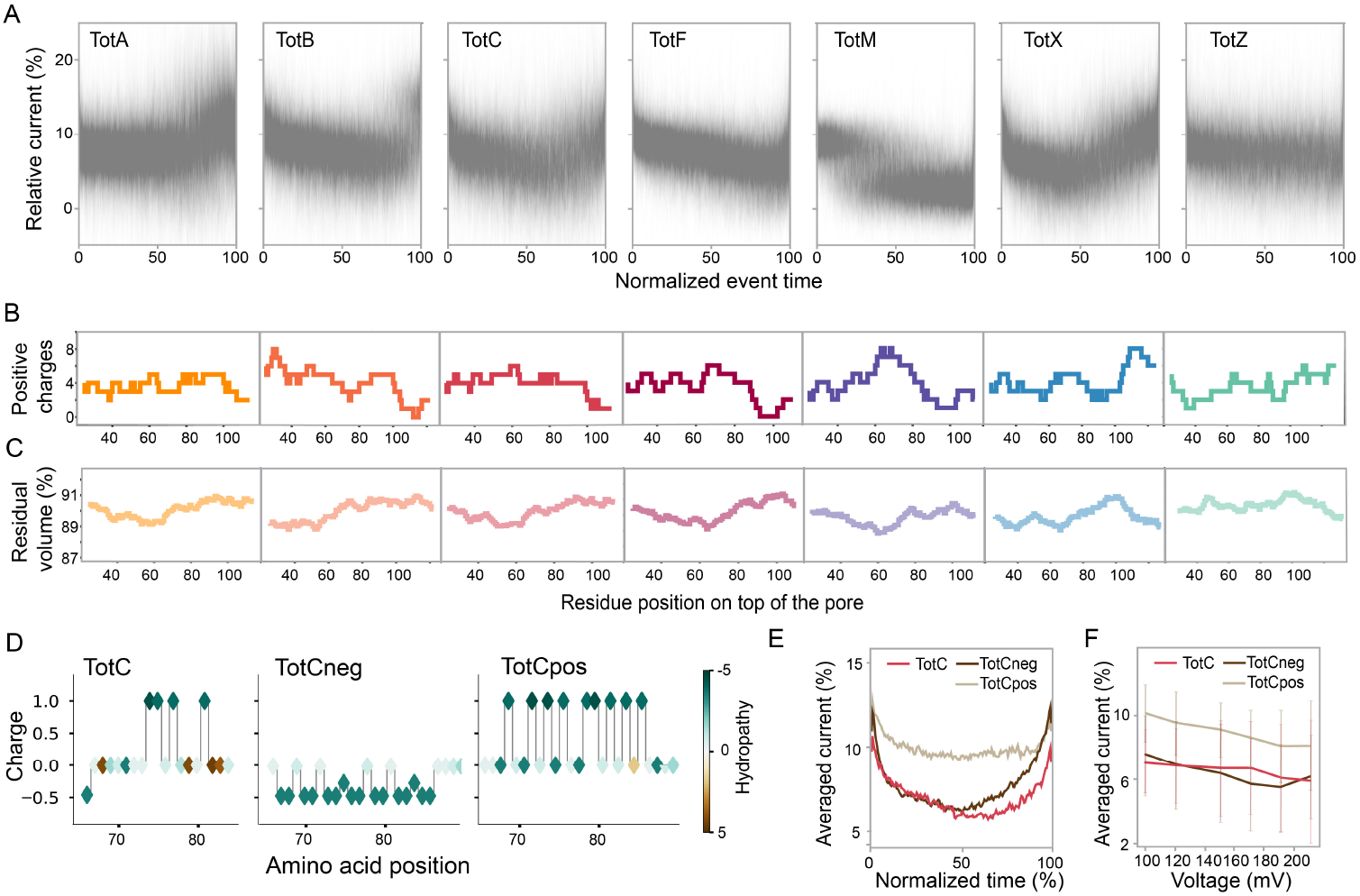
Interpretation of current signals. (A) Time normalized overlay plots of 800 randomly chosen events for each Tot family protein recorded at 100 mV. Sum of positive charges (B) and relative residual volume (C) for a sliding window of 30 amino acid along the sequence of TotA (yellow), TotB (orange), TotC (light red), TotF (dark red), TotM (purple), TotX (blue), TotZ (turquoise). (D) Charge property along the unstructured loop of TotC, TotCneg, and TotCpos (left to right). (E) Average current of 800 randomly chosen events along normalized time recorded at 100 mV for TotC, TotCneg and TotCpos. (F) Population average over the mean currents of TotC (light red), TotCneg (brown), and TotCpos (green). Overlay plots of each protein are shown in SI Fig. S12.

To interpret the current signatures, we simplified the system by assuming that the simultaneous effect of all residues inside the pore lumen results in one current level. We considered a sliding window of size *l* = 30 aa – approximately corresponding to the pore height – gliding along the analytes and monitored the sum of positive charge in each window (Fig. 4B, SI Fig. S14). Strikingly, key features of the overlayed currents are recovered in these sliding windows of positive charges. The sliding windows represent N-terminal capture, since calculations from C-terminus capture showed lower agreement with experimental data. We further verified the effect of positive charges by replacing the unstructured loop of TotC with two control sequences (Fig. 4C): one with more negative charges (TotCneg), and one with more positive charges (TotCpos), their properties are shown in Fig. 4D. Indeed, when looking at the event patterns we found a higher current level in the signals for TotCpos, which was not present with TotC (Fig. 4E). Additionally, the population average for the current of TotCpos translocations (SI Fig. S15) was higher compared to TotC and TotCneg over all voltages (Fig. 4F). TotCneg behaved very similarly to TotC, suggesting that the positive charges produced a much more pronounced effect than negative ones in our system. This can be explained by the asymmetry of the ion transport in the GdmCl pH 4.0 system, where anions are more mobile than cations. Introducing higher positive charge inside the pore can thereby increase anion (Cl^-^) fluxes and the total current. A similar effect was observed in the open pore current, which increased when the pore lumen was more positively charged at pH 4.0 compared to pH 7.5 (SI Fig. S4). As negative charged aa occupy the pore, they neutralize part of the asymmetry in ion transport and do not lead to a current increase.

A typical model to explain nanopore sensing is volume exclusion^43^. In studies examining how aa properties influence current readouts – for instance when protein analyte motion through MspA or CsgG was controlled using a Helicase^14,44^ or ClpX^17^ – the volume of the amino acids within the reading site often correlated with the observed current blockages. To investigate volume exclusion in our system, we estimated the volume of the aerolysin pore lumen as 41.5 nm^3^ and considered the relative residual volume during protein translocations using 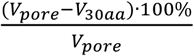, where V30aa is the summed volume of aa^45^ in each sliding window (see methods for details). As seen in Fig 4C, the relative residual volume is similar across our different analytes, and no correlation can be drawn between the volume and the event patterns. This could be because of the free translocation approach we investigated here. While a large part of the current blockage undoubtedly stems from volume exclusion, we expect the current to be influenced by an average over 30 aa and therefore the differences in volume exclusion are likely not substantial enough to significantly influence the observed signal patterns.

## Discussion

As the field of nanopore protein analysis evolves, the technology is poised to become indispensable, offering speed, affordability, portability, and throughput compared to traditional methods. In this study, we investigated the capabilities of an aerolysin nanopore for label-free fingerprinting of full-length proteins. Our analysis underscores the critical role of the EOF as a driving force in protein measurements. While the long and narrow barrel of aerolysin provides a big surface that can yield large ion selectivity, it also provides more hydrodynamic resistance compared to other nanopores which reduces the resulting water flux^28^. This explains why in 2 M GdmCl at pH 7.5, TotA cannot translocate through aerolysin, whereas other proteins have been translocated through α-HL under the same condition^22^. Here, we overcome this challenge by combining GdmCl with a low pH condition which induces an even larger EOF. We observed that the residual structure of TotA at 2 M GdmCl did not impact its translocation or signal patterns. This is an important advantage, as even at 3 M GdmCl, some protein structures may remain. Generating reliable signals under such conditions is crucial for a broad applicability in protein identification.

By evaluating related proteins, we observed that translocation kinetics were strongly influenced by interactions between sequence-specific properties of the protein and the pore. Aerolysin K238A demonstrated promising potential for protein identification when coupled with machine learning algorithms. By optimizing the applied voltage, we achieved 80% classification accuracy for seven proteins from the same family, which shared 38%–70% pairwise sequence identities. This highlights the potential of the nanopore system for high-accuracy, label-free protein identification. While a fingerprinting approach cannot directly provide the analytes sequence, its ability to operate without enzymatic processing or chemical labeling, combined with machine learning-based classification, positions it as a practical and efficient tool.

Our findings suggest that the sequence-specific current patterns observed in aerolysin are predominantly influenced by the distribution of positive charges along the protein sequence.

Positive charges of analyte proteins within our nanopore system appear to increase the measured current by elevating chloride ion fluxes. This model provides a rational explanation for the observed signal patterns and offers a foundation for future advancements. With continued development, like identifying additional factors that influence the current output and translocation kinetics, or engineer/design pore architecture to accommodate less amino acid and therefore higher resolution, it may become feasible to predict nanopore fingerprints from protein sequences or, conversely, infer *de novo* sequence information directly from single-molecule signals, shedding light on previously uncharted biology.

## Acknowledgements

We thank Dr. Samuel Rommelaere and Prof. Bruno Lemaitre for the generous gift of the proteins TotA, TotB, TotC, TotF, TotM, TotX, and TotZ. This work was supported in part using the resources and services of the Protein Production and Structure Core Facility at the School of Life Sciences of EPFL. We thank Dr. Kelvin Lau for support and scientific discussion concerning protein production. This work was supported by Swiss National Science Foundation (PR00P3_193090 to C.C).

## Author contributions

V.R. and C.C. conceived the project and designed the study. V.R. performed circular dichroism experiments. E.N. performed reversal potential measurements and analysis. V.R., E.N., G.B., and J.G. performed sensing experiments and signal processing. V.R. analyzed the data and performed machine learning classifications. J.D. performed molecular dynamics simulations and analysis. V.R. and C.C. interpreted the data. V.R., and C.C. wrote the manuscript with input from all authors.

