## SupplementaryInformation for "Charge-based fingerprinting of unlabeled full-length proteins using an aerolysin nanopore"

### Methods

#### Materials

Aerolysin K238A was produced and activated as described previously[1]. Turandot protein sequences were cloned into pET29b vectors, which were ordered from GenScript. Production was performed as described previously[2] by the Protein Production and Structure Core Facility at EPFL. TotA, TotB, TotC, TotX, TotZ, TotCpos, and TotCneg were expressed with a C-terminal His tag, while TotM and TotF were produced with C-terminal MBP-His constructs. A TEV cleavage site was included to allow removal of either tagging strategy. The plasmid encoding the protein of interest was transformed into Escherichia coli BL21 (DE3) cells. A 1L culture of Terrific Broth (TB) supplemented with kanamycin was inoculated and grown at 37°C until the optical density at 600 nm (OD600) exceeded 0.6. Protein expression was induced with 0.5 mM isopropyl β-D-1-thiogalactopyranoside (IPTG), and the culture was incubated at 18°C overnight. Two main buffers were used for the purification process: Buffer A – 700 mM NaCl, 20 mM HEPES, pH 7.5.; Buffer B – 700 mM NaCl, 20 mM HEPES, pH 7.5, 500 mM Imidazole. For dialysis, a buffer containing 20 mM HEPES, pH 7.5, and 300 mM NaCl was used. The bacterial pellet was resuspended and lysed. The supernatant was incubated with 5 mL of HisPur Ni-NTA Resin beads (Thermo Scientific) at 4°C for 2 hours and eluted on a gradient from buffer A to 100% buffer B. Fractions containing the protein of interest were pooled and subjected to overnight cleavage with 0.7 mg SuperTEV protease at 4°C during dialysis. Cleaved proteins underwent reverse Ni-NTA chromatography to remove the protease and cleaved tag. The purified and concentrated protein samples were flash-frozen and stored at -80°C for future use.

#### Nanopore sensing experiments

Nanopore measurements were carried out in the indicated buffer conditions containing 2 M GdmCl (Sigma) or 3 M GdmCl buffered either with 10 mM tris(hydroxymethyl)aminomethane to pH 7.5 or with 10 mM citrate to pH 4.0. Lipid bilayers of 1,2-diphytanoyl-sn-glycero-3-phosphocholine (DPhPC, Avanti polar Lipids) were formed on the 50 µm apertures of MECA 4 Recording Chips (Ionera, Germany) and small amounts of activated aerolysin K238A pore were added to the *cis* chamber to allow single pore insertion. Data was acquired at 100 kHz using an Orbit mini device (Nanion technologies, Germany). Temperature control of the instrument was set to 25°C for all measurements. Each experiment was carried out several times and with > 5 individual pores. During condition optimization, TotA was measured at 0.5 µM initially. All subsequent measurements were carried out at 1 µM analyte concentration.

Reversal potential measurements

To obtain the reversal potential, measurements were carried out on Axonpatch 200B (Molecular Devices) using polystyrene cups (Warner Instruments). 1 µl of 25 mg/ml DPhPC in hexane (Sigma) was used to pretreat the cup. Following evaporation, 1225 µl of 0.5 M electrolyte were added to both the *cis* and *trans* chamber. Electrodes were installed with salt bridges and membranes were formed using 8 mg/ml DPhPC in decane (Sigma). Small amounts of activated aerolysin K238A were added to the *cis* chamber. Following a pore insertion, the pipette offset was used to ensure that 0 pA were measured at 0 mV in symmetric buffer conditions (0.5 M KCl). Subsequently, 525 µl of the *cis* buffer were exchanged for buffer at the same pH with 4 M electrolyte for a final 2 M in *cis* versus 0.5 M in *trans*. Following carful mixing and several minutes of equilibration time, the voltage was altered in 2 mV steps within the range of ± 40 mV. As the current depended linearly on the voltage in this regime, a linear regression allowed to precisely extract the voltage at 0 pA, which is known as reversal potential ($V_R$). Activity coefficients[3,4] were used to convert concentrations to activity $a$. The reversal potential $V_R$ was used to calculate $P^-/P^+$ with the Goldman–Hodgkin–Katz flux equation (Equ. 1) [5].

$$\frac{P^-}{P^+} = \frac{[a_{cation}]_{cis} \cdot \exp\left(-\frac{eV_R}{k_B T}\right) - [a_{cation}]_{trans}}{[a_{anion}]_{cis} - [a_{anion}]_{trans} \cdot \exp\left(-\frac{eV_R}{k_B T}\right)} \qquad \text{Equ. 1}$$

With the Boltzmann constant $k_B$, elementary charge $e$ and the absolute temperature $T$. Final values are averaged over results from at least 4 individual pores (Fig. S1).

Pretreatment of protein analytes

Samples were equilibrated in 6 M GdmCl and heated to 95°C for 10 min before being measured in the nanopore system or in Circular Dichroism.

Circular dichroism

Circular dichroism (CD) measurements were performed using a Chirascan™ V100 (Applied Photophysics) spectropolarimeter. Samples were analyzed in quartz cuvettes with a path length of 0.1 cm. Spectra were recorded from 210 to 280 nm at 25°C. Each spectrum was an average of 3 accumulations. The air baseline was automatically subtracted from all measurements by the instrument. Buffer spectra were recorded separately and used as backgrounds to be subtracted for each corresponding sample. TotA was measured following the pre-equilibration protocol at 10 µM final concentration in all conditions. Data processing and analysis were performed in python. The results are presented as mean residue ellipticity ($deg \cdot cm^2 \cdot dmol^{-1}$) with consideration of protein concentration, path length, and number of amino acid residues.

Data analysis

The event-current representing single molecule interactions was extracted from raw traces in python. Briefly, the traces were filtered to 15 kHz with an eighth order low pass Bessel filter and segmented at each deviation in voltage. For each segment, the open pore current (OPC, $I_0$) and its standard deviation (sigma) were determined. Regions in the trace that correspond to events were defined as starting when the current dropped below 10 sigma from the OPC and ending when the current had recovered to 1 sigma below OPC. Events were extracted from the unfiltered raw traces using the regions determined in filtered traces. The gradient of the datapoints was thresholded at the beginning and end of the event to exclude datapoints from the falling and rising regions of the measured current.

To exclude background and outlier data, events with less than 15% mean relative residual current and dwell time of 0.25 – 100000 ms were selected and further cleaned using the Isolation Forest algorithm from the Scikit-learn library. For this, the mean, mode, and median of the relative current, as well as its standard deviation and the logarithmic dwell time of the event were used as event features. Dwell times of the populations were fitted with an exponentially modified gaussian distribution or the Fokker-Planck equation[6] (Equ. 2) with *L* the contour length of each protein plus the length of the pore (estimated at 10 nm), the dwell times *t* , the diffusion coefficient *D*, and the drift velocity *v*.

$$F(t) = \frac{L}{\sqrt{3\pi D t^3}} \cdot \exp\left(-\frac{(b - vt)^2}{4Dt}\right) \qquad \text{Equ. 2}$$

Since the equation overly simplifies the system, fitted *D* and *v* values were not considered reliable and are thus not discussed. The equation was used merely to extract each distributions' maximum and half widths.

Time normalized overlay plots were produced as done in our previous study[7]. Briefly, currents ($I/I_0$ in %) of randomly selected events were normalized to a length of 200 data points by dividing them into 200 equal chunks and averaging them. In cases of events with less than 200 data points, they were interpolated using numpy's interp function.

Machine learning

For identification, additional features were extracted from the event data (listed in SI Table S2). The current of each event was Bessel filtered to 10 kHz and divided into 8 equally long parts, or three equally long parts as indicated in SI Table S2. Features were also extracted from the power spectrum of each event by segmenting the spectrum into 8 parts and averaging it similar as in a previous study[8]. Additionally current values were separated into two clusters using a gaussian mixture model and averaged, yielding a low current and high current average for each event.

The maximum equal number of events were used to fit a random forest classifier from the sikit-learn library and only features with a mean decrease in impurity of over 0.01 were selected for further classification (SI Fig. S11). Classification algorithms from the sikit-learn library were cross validated and the top three performing classifiers (Random Forest, Bagging, and Extra Trees Classifier) were used in a voting classifier. To estimate the quality of protein identification, accuracy was estimated in 10 rounds and averaged. For each round, the equal number of events was randomly sampled from each protein analyte, followed by a split that left 20% of the data points for testing. A pipeline of standard scaler and voting classifier was built and trained on the training data set.

<u>All-atom MD simulation</u>

*Building MD simulation system*

We prepared the molecular dynamics (MD) simulation systems using the cryo-EM structure of wild-type aerolysin (PDB: 9FM6)[1]. CHARMM-GUI was used to generate the mutant K238A, configuration, and topology of the simulation systems[2]. The parameter files were created using the CHARMM36 force field[3]. We used the KBFF20 parameter for $Gdm^+$ ion[4]. The PPM server was used to reorient the aerolysin structure, ensuring that its transmembrane region was correctly positioned in a lipid bilayer[5]. The protein was embedded in a PhPC bilayer. Next, we solvated the protein-membrane complex in a water box using the TIP3 water model. Cations and anions were added to achieve a concentration of 3 M while counterions were introduced to neutralize the system. Further details of simulation systems are shown in SI Table S1.

All of the MD simulations were performed with GROMACS 2023.1[6]. The REDUCE program in AMBER was used to add hydrogens to the original PDB files and determine the protonation state at pH 7.5[7]. To accurately calculate the protonation state at low pH, we used the APBS server for pH 4.0 systems to correct the protonation state[8].

We used the steepest descent algorithm to achieve energy minimization. Then, following up with a two-stage equilibration, a 0.4 ns NVT equilibration simulation with harmonic restraint was applied to the protein molecule (force constants of 4000 $kJ \cdot mol^{-1} \cdot nm^{-2}$ on the backbone and 2000 $kJ \cdot mol^{-1} \cdot nm^{-2}$ on the side chains), and a 20 ns NPT equilibration simulation with gradually decreased restraint (from 2000 to 100 $kJ \cdot mol^{-1} \cdot nm^{-2}$ on the backbone and from 1000 to 50 $kJ \cdot mol^{-1} \cdot nm^{-2}$ on the side chains). During the equilibration processes, planar restraints were used to keep the positions of lipid head groups along the membrane-normal direction.

The simulation temperature of the system was set to 300 K. The time step was 2 fs. The cubic periodic boundary condition was used during the simulations, and the van der Waals interaction was switched off from 10 to 12 Å. The long-range electrostatic interactions were calculated using the particle mesh Ewald (PME) method[9].

To investigate the ion transport properties of the pore and understand its corresponding ion selectivity feature, we applied a 100 mV transmembrane potential to the channel and analyzed the ion translocation process across the pore.

*Analysis of electrostatic potential map*

We calculated the surface electrostatic potential of the protein with APBS electrostatic plugin in VMD[8].

*Analysis of the binding distribution of ions*

We used the HOLE program to get the inner radius of the aerolysin K238A structure[10].To analyze the binding of ions to the pore lumen we only consider the ions that enter the protein channel within the trajectory. For each ion type and position along the pore lumen we then sum the time during which binding occurred. Binding was identified, when the distance between the central carbon atom of a $Gdm^+$ ion – or for other ions the exact position of the atom – and the pore wall was less than 4 Å. This is similar to what has been defined previously[11]. For the ions that entered the pore, the ratio of each bound ion was calculated by dividing the accumulated binding time along the Z coordinate by the total time of the trajectory. Final distributions along the Z coordinate were averaged over the ions that entered the pore.

### Current interpretation

To interpret current signatures, each amino acid in a sequence was first replaced by its charge or volume [20]. Charges were calculated based on the pH and pKa[21] of the residues and terminal groups. Terminal charges were added to the first and last amino acid and all negative values were set to 0 to only consider the positive charges. The sum of either positive charges or volume of each set of 30 subsequent amino acids along the sequence was calculated. This was done in direction from N- to C- terminus, since the signal features of all analytes were better compatible with N-terminal capture. The summed volume of the analyte portion ($V_{30aa}$) was further converted into relative residual volume as $\frac{(V_{pore}-V_{30aa})\cdot 100\%}{V_{pore}}$ using the approximate pore volume $V_{pore} = \pi r^2 h$. The pore volume was calculated using the average water accessible radius of the pore ($r = 1.15 \text{ nm}$) and the height of the aerolysin barrel ($h = 10 \text{ nm}$).

**SI Note**

Translocation velocities (Fig. 3E, proteins length divided by the fitted dwell time) depended roughly linearly on the electric field $E = \frac{\Delta V}{d}$, with the voltage drop $\Delta V$ over the length of the pore ($d \approx$ 10 nm). The transport in this system is driven by the EOF against the EPF and we therefore observe $v_{total} = v_{EOF} + v_{EP}$. The electroosmotic flow velocity $v_{EOF}$ in biological nanopores is directly proportional to the current $I$[37], which in turn depends linearly on the voltage in the measured regime (SI Fig. S4). The electrophoretic velocity has been described as $v_{EP} = \mu_{EP} \cdot E = \mu_{EP} \cdot \frac{\Delta V}{d}$ with the electrophoretic mobility $\mu_{EP}$. Thus, the observed linear dependence of the velocity on the electric field is in alignment with expectations.

### SI Tables

Table S1. MD simulation systems of aerolysin K238A in different buffer and pH.

| Trajectory label | pH | Ions | Box size (Å) | Atom number | Trajectory time (ns) |
|---|---|---|---|---|---|
| Traj-1 | 4.0 | 3 M GdmCl | 199×199×150 | 610,531 | 450 |
| Traj-2 | 7.5 | 3 M GdmCl | 198×198×150 | 611,415 | 450 |
| Traj-3 | 4.0 | 3 M KCl | 195×195×151 | 583,387 | 300 |
| Traj-4 | 7.5 | 3 M KCl | 197×197×149 | 573,545 | 200 |

Table S2. event features used for machine learning classification. Except for feature 0 (dwell time) all features are described as calculations on the extracted current that was normalized by the open pore current and filtered to 10 kHz. For features 7 and 8 current data point were divided into two clusters using a gaussian mixture model.

| | | | | | |
|---|---|---|---|---|---|
| 0 | logarithmized dwell time | 31 | standard deviation part 7/8 | 62 | max part 2/8 |
| 1 | mean | 32 | standard deviation part 8/8 | 63 | max part 3/8 |
| 2 | mode | 33 | mean power spectrum part 1/8 | 64 | max part 4/8 |
| 3 | median | 34 | mean power spectrum part 2/8 | 65 | max part 5/8 |
| 4 | standard deviation | 35 | mean power spectrum part 3/8 | 66 | max part 6/8 |
| 5 | skewness | 36 | mean power spectrum part 4/8 | 67 | max part 7/8 |
| 6 | kurtosis | 37 | mean power spectrum part 5/8 | 68 | max part 8/8 |
| 7 | mean of lower current cluster | 38 | mean power spectrum part 6/8 | 69 | skewness part 1/8 |
| 8 | mean of higher current cluster | 39 | mean power spectrum part 7/8 | 70 | skewness part 2/8 |
| 9 | median part 1/8 | 40 | mean power spectrum part 8/8 | 71 | skewness part 3/8 |
| 10 | median part 2/8 | 41 | mean absolute gradient | 72 | skewness part 4/8 |
| 11 | median part 3/8 | 42 | mean gradient | 73 | skewness part 5/8 |
| 12 | median part 4/8 | 43 | mean abs gradient 1/3 | 74 | skewness part 6/8 |
| 13 | median part 5/8 | 44 | mean abs gradient 2/3 | 75 | skewness part 7/8 |
| 14 | median part 6/8 | 45 | mean abs gradient 3/3 | 76 | skewness part 8/8 |
| 15 | median part 7/8 | 46 | diff mean 12 | 77 | mode part 1/8 |
| 16 | median part 8/8 | 47 | diff mean 23 | 78 | mode part 2/8 |
| 17 | mean part 1/8 | 48 | diff mean 34 | 79 | mode part 3/8 |
| 18 | mean part 2/8 | 49 | diff mean 45 | 80 | mode part 4/8 |
| 19 | mean part 3/8 | 50 | diff mean 56 | 81 | mode part 5/8 |
| 20 | mean part 4/8 | 51 | diff mean 67 | 82 | mode part 6/8 |
| 21 | mean part 5/8 | 52 | diff mean 78 | 83 | mode part 7/8 |
| 22 | mean part 6/8 | 53 | min part 1/8 | 84 | mode part 8/8 |
| 23 | mean part 7/8 | 54 | min part 2/8 | 85 | correlation part 1/3 |
| 24 | mean part 8/8 | 55 | min part 3/8 | 86 | correlation part 2/3 |
| 25 | standard deviation part 1/8 | 56 | min part 4/8 | 87 | correlation part 3/3 |
| 26 | standard deviation part 2/8 | 57 | min part 5/8 | | |
| 27 | standard deviation part 3/8 | 58 | min part 6/8 | | |
| 28 | standard deviation part 4/8 | 59 | min part 7/8 | | |
| 29 | standard deviation part 5/8 | 60 | min part 8/8 | | |
| 30 | standard deviation part 6/8 | 61 | max part 1/8 | | |

### SI Figures

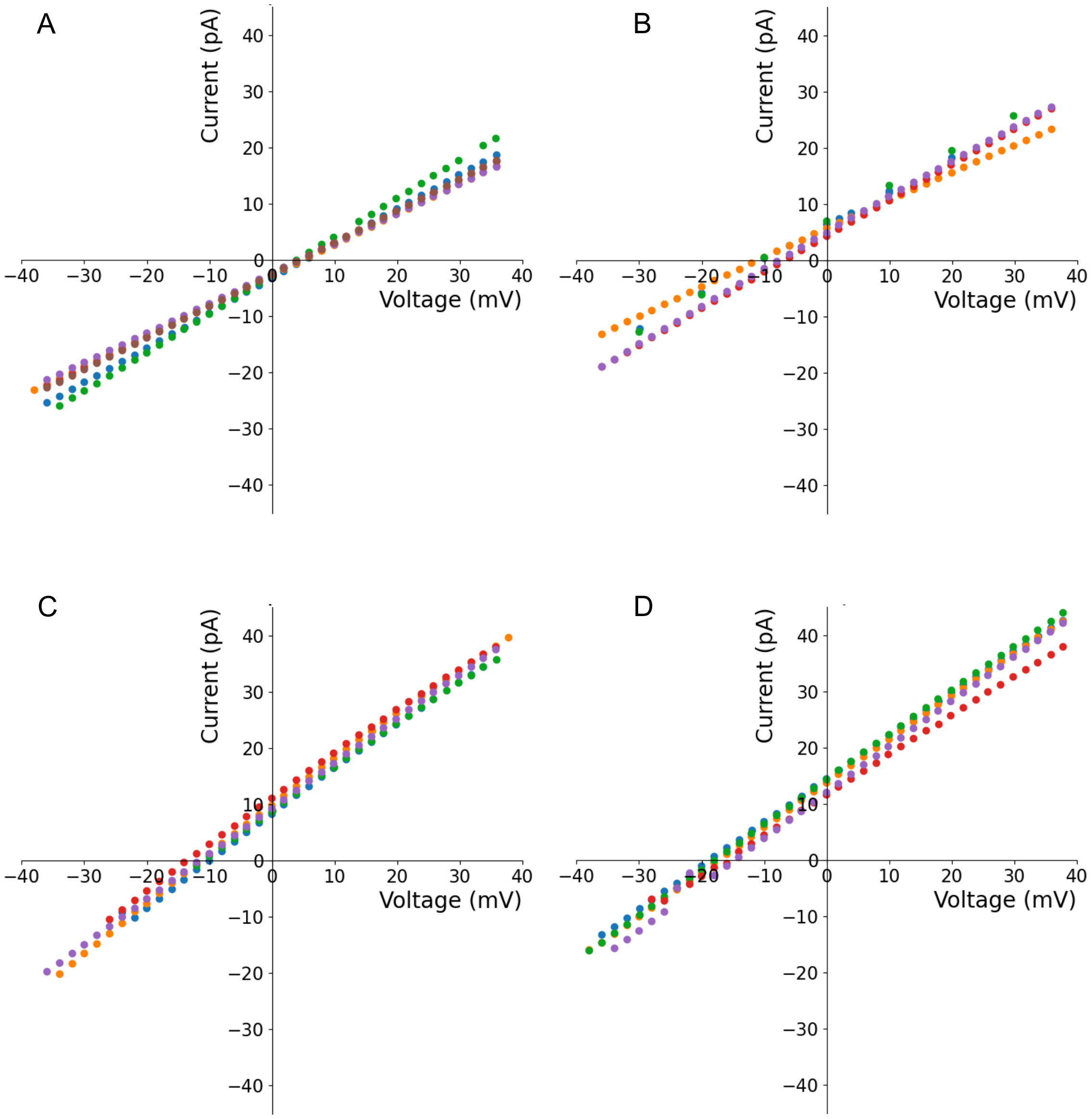


Figure S1. Current versus voltage data at 2 M *(cis)* vs 0.5 M *(trans)* electrolyte measured with aerolysin K238A *in* KCl at pH 7.5 (A), GdmCl at pH 7.5 (B), KCl at pH 4.0 (C), and GdmCl at pH 4.0.

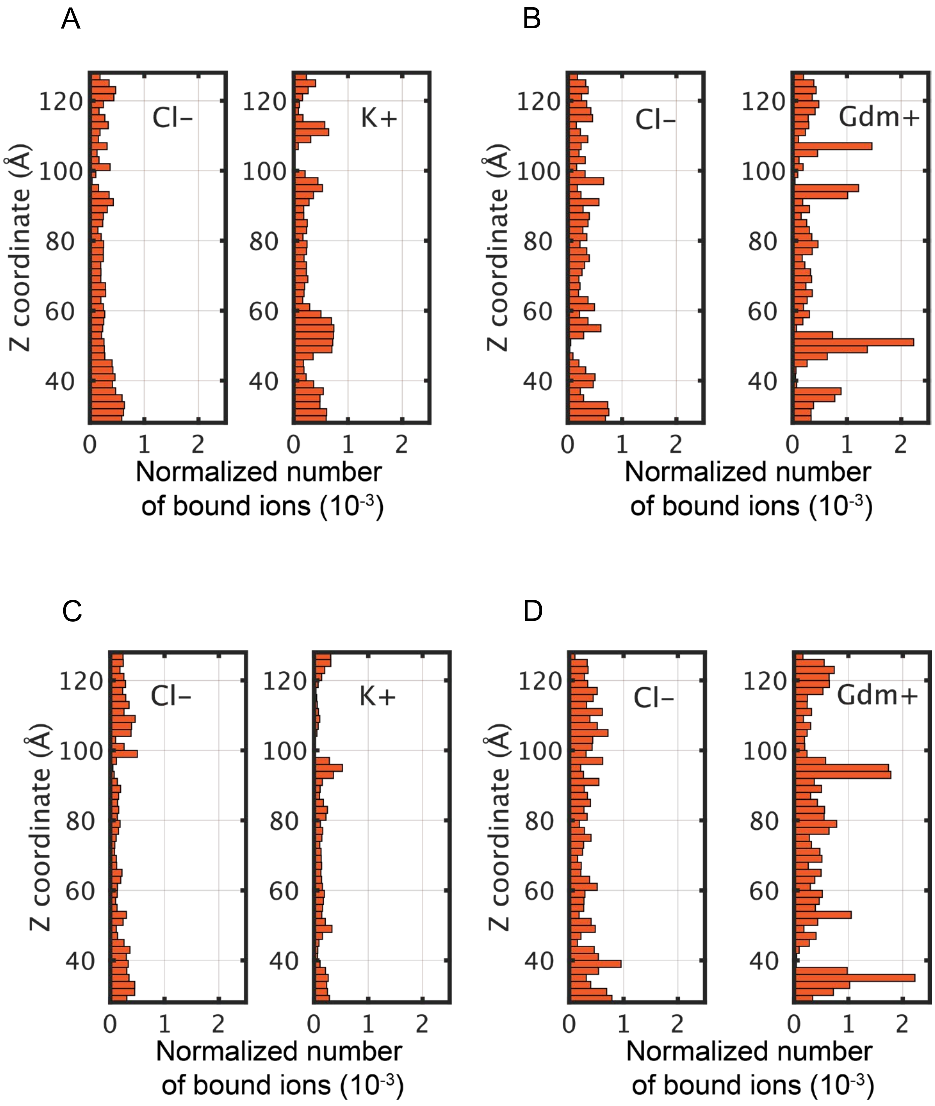


Figure S2. MD results displaying the duration of ions binding to the surface of aerolysin K238A normalized by the number of interacting ions. (A) 3M KCl at pH 7.5, (B) 3M GdmCl at pH 7.5, (C) 3M KCl at pH 4.0, (D) 3M GdmCl at pH 4.0.

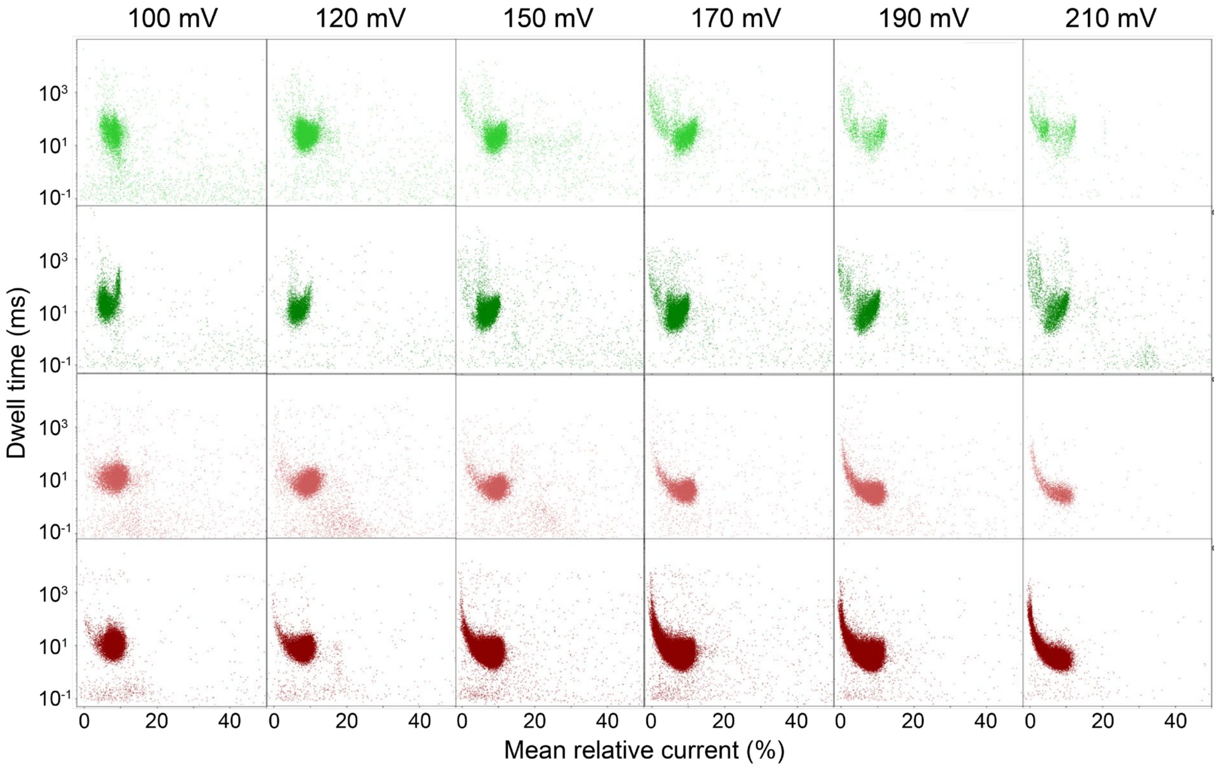


Figure S3. Scatter plots of individual single events from TotA represented by event duration against of mean relative current. From top to bottom: 2 M GdmCl pH 7.5, 3 M GdmCl pH 7.5, 2 M GdmCl pH 4.0, 3 M GdmCl pH 4.0 at (left to right) 100 mV, 120 mV, 150 mV, 170 mV, 190 mV, 210 mV.

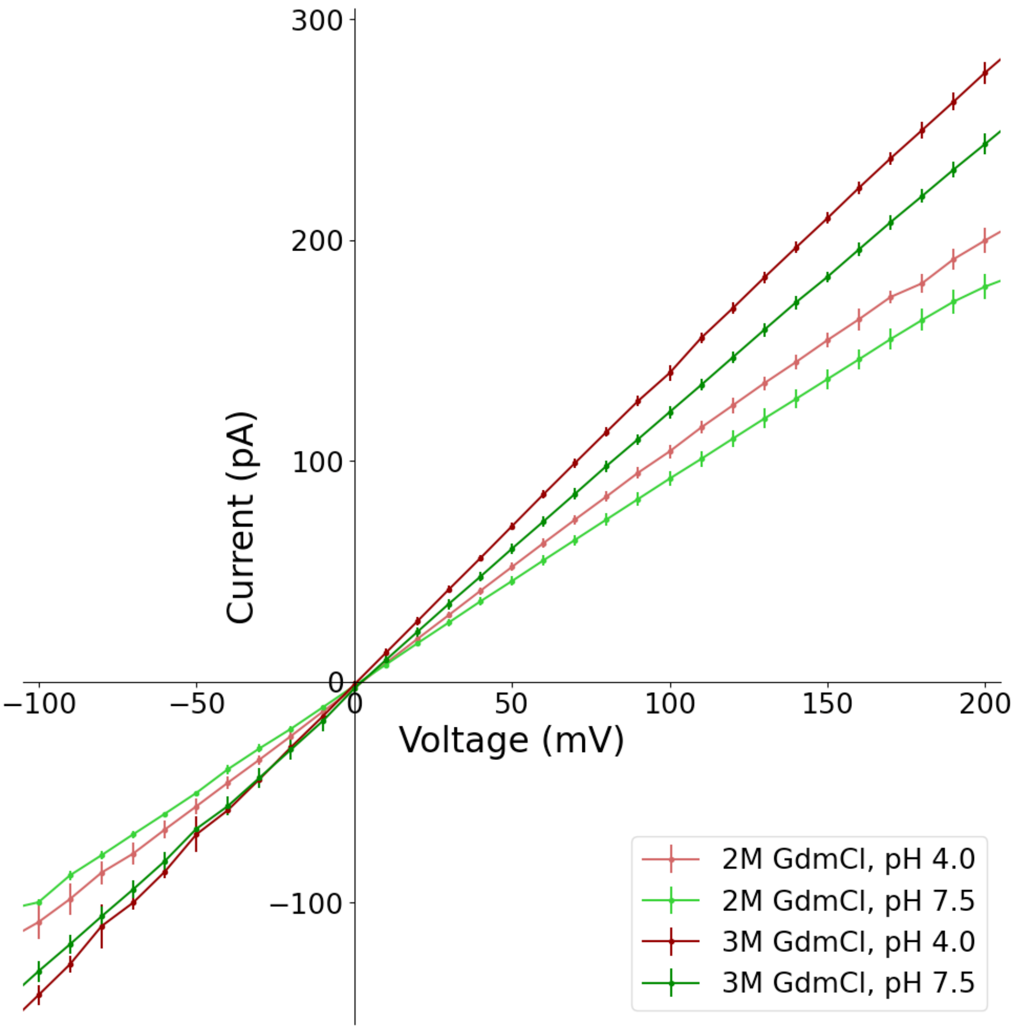


Figure S 4. Open pore current of aerolysin K238A at various voltages in the indicated conditions.

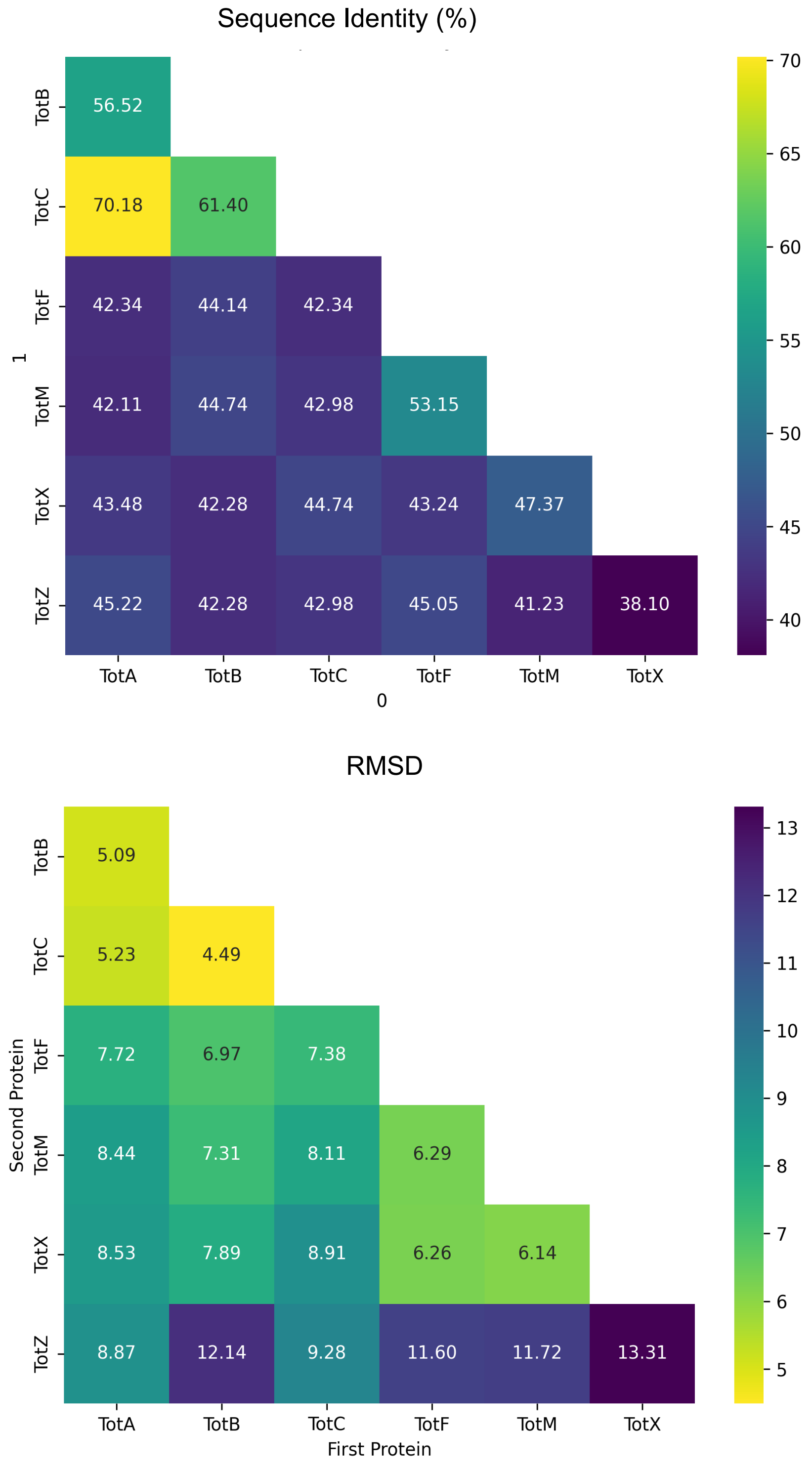


Figure S5. Sequence similarity (left) and structural similarities (right) calculated by Root mean square deviation (RMSD) of Tot family proteins

TotA:

MGYSDEDREADNLRIAEIIKNAQDDDSKINSTQELLDIYRRLYPSLTPEERESIDKFVNEHTDAIIID
GVPIQGGRKARIVGKIVSPGVKGLATGFFEELGSKLAQLFAGENLYF

TotB:

MYSNQERQRDSRRVAEIMRTSWDDNTKIKRIQELLTIYNRMAPSLRPDERARMDRFISGYTGEIM
VDGVPSQGGARRIFKKILSPAAKSVATGFFTELGASLASILTSWFPANTERNHENLYF

TotC:

MYSDEERESDSLRVAEIIRTSNDAESKINRTQELLDIFRRLTPTLSPEQREKIERSIQEHTDEILIDG
VPSQGGRKTKYVGKILSPVAQGLAVGFFEELGGSLSRLFTGENLYF

TotF:

MEHAQSDPEFTAKARQMLAVFGNSEVDRYTKSRNLPALIEFYEKYSSRLPLTVQDRTYANNVIRR
YRAHNNQQVDGVPAQGGVGVVFALLLPFAVSIVEGIAKAIRENLYF

TotM:

MENEDEFVTEKQRLFSVYGDSSVDEATKYRNIDSLVTFYDKYFTRLQLKPDLNTRAHDLLRRYKE
ENARVVLVDGTPAQGGFWLPLVKLLIVQLGVEIASEGVKRAIESENLYF

TotX:

MNTNSSSYEEHRNYLLNIFHNPFVNDSIKEKNIPQLIAFYQRYPTDVPLSDADRQQFERFIHDYRE
YRAVLVDGAPPQGGSFGNIFGHFLGRVGTRYISSLFNKKREERKSNHAYIIEDYNENLYF

TotZ:

MRMLDADRNRLQQLQIRSQQSADANTQVDIAYEVIGIYDKYKGQGGSNVLREAQLNSQVNDFK
RKTMVIDGVPAQGGVWGILGAIKKAADAVPDNVKKDAENLVKSSTKVLVRGIYDYLMGKMKHEN
LYF

TotCpos:

MYSDEERESDSLRVAEIIRTSNDAESKINRTQELLDIFRRLTPTLSPEQREKIERSIQEHTDEILIGG
NKSNRSRNKGQKRSKSKAKGQSPVAQGLAVGFFEELGGSLSRLFTGENLYF

TotCneg:

MYSDEERESDSLRVAEIIRTSNDAESKINRTQELLDIFRRLTPTLSPEQREKIERSIQEHTDEILIGD
DSDDGDDEDDSDDGDDEDDGGSPVAQGLAVGFFEELGGSLSRLFTGENLYF

Figure S6. Protein sequences of measured analytes.

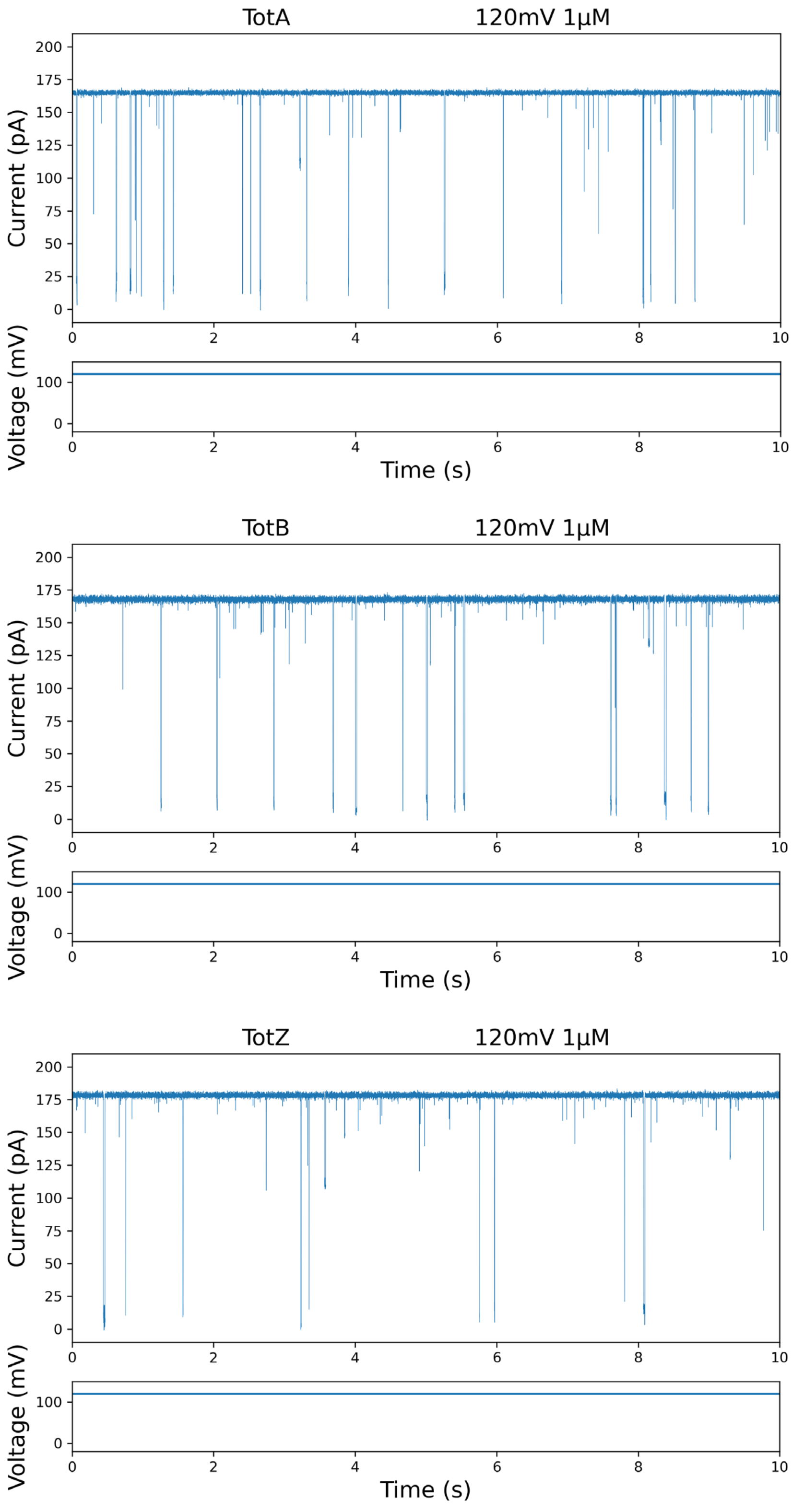


Figure S7. Example traces of TotA, TotB, and TotZ recorded in 3 M GdmCl at pH 4.0, 120 mV, filtered with 1 kHz for illustration purpose only.

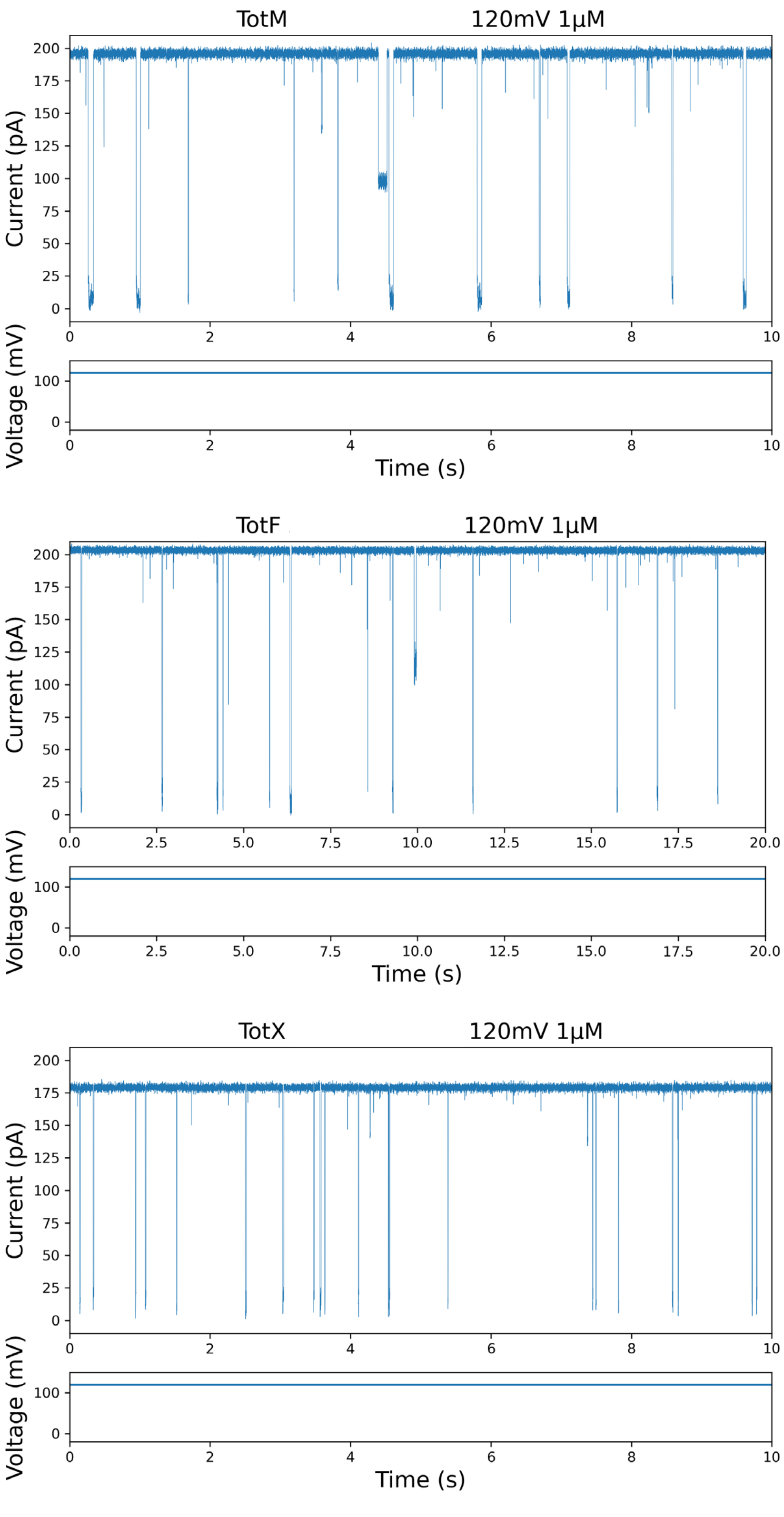


Figure S8. Example traces of TotF, TotM, and TotX recorded in 3 M GdmCl at pH 4.0, 120 mV, filtered with 1 kHz for illustration purpose only.

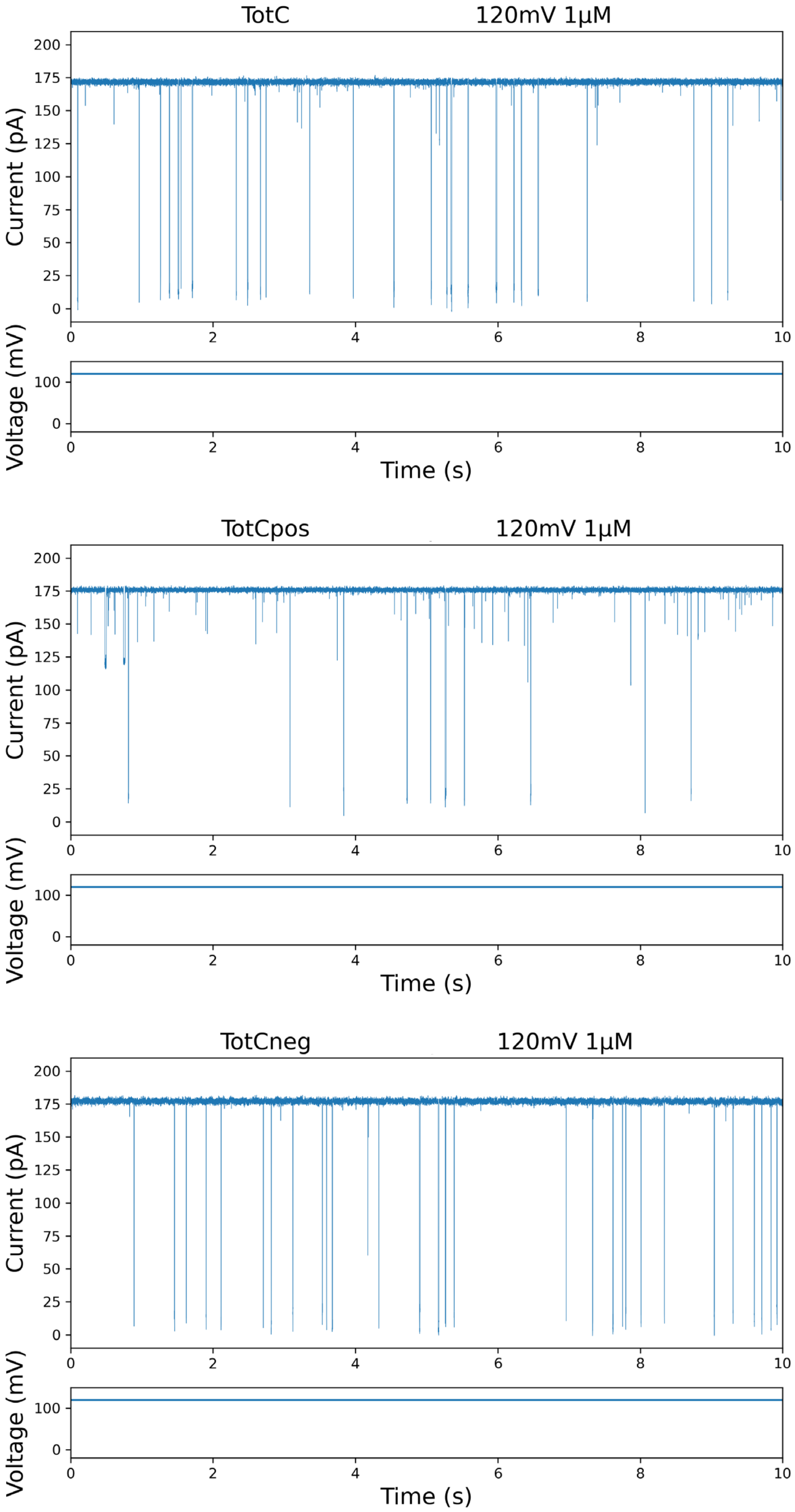


Figure S9. Example traces of TotC, TotCpos, and TotCneg recorded in 3 M GdmCl at pH 4.0, 120 mV, filtered with 1 kHz for illustration purpose only.

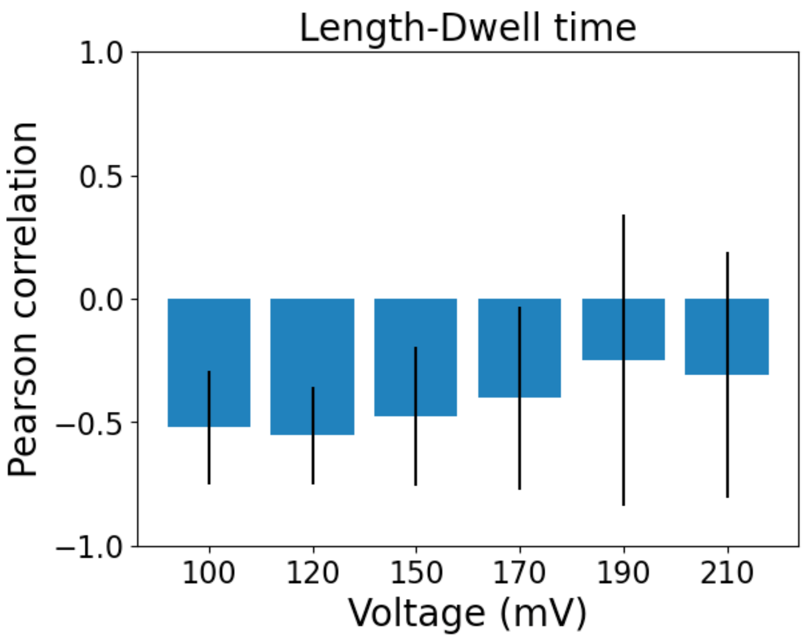


Figure S10. Pearson correlation between analyte length (number of amino acids) and the fitted dwell time for the 7 Tot family proteins across different voltages.

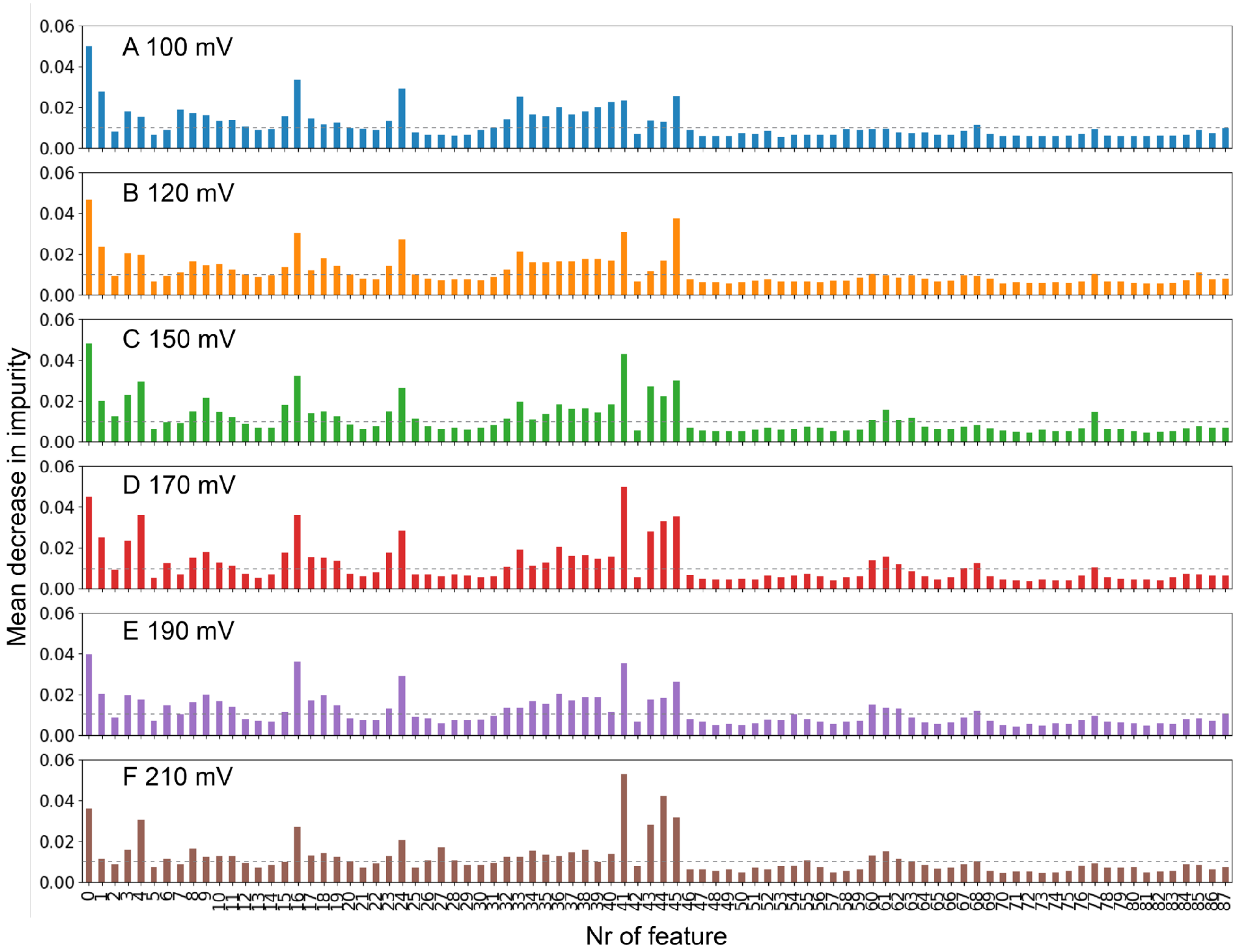


Figure S11. Mean decreases of impurity for 7-way classifications of TotA, TotB, TotC, TotF, TotM, TotX, and TotZ in 3 M GdmCl at pH 4.0 measured with aerolysin K238A at (A) 100 mV, (B) 120 mV, (C) 150 mV, (D) 170 mV, (E) 190 mV, and (F) 210 mV. The feature number corresponds to features listed in SI Table S2. In classifications only features that exceeded the threshold of 0.1 MDI (shown as grey, dashed line) were used.

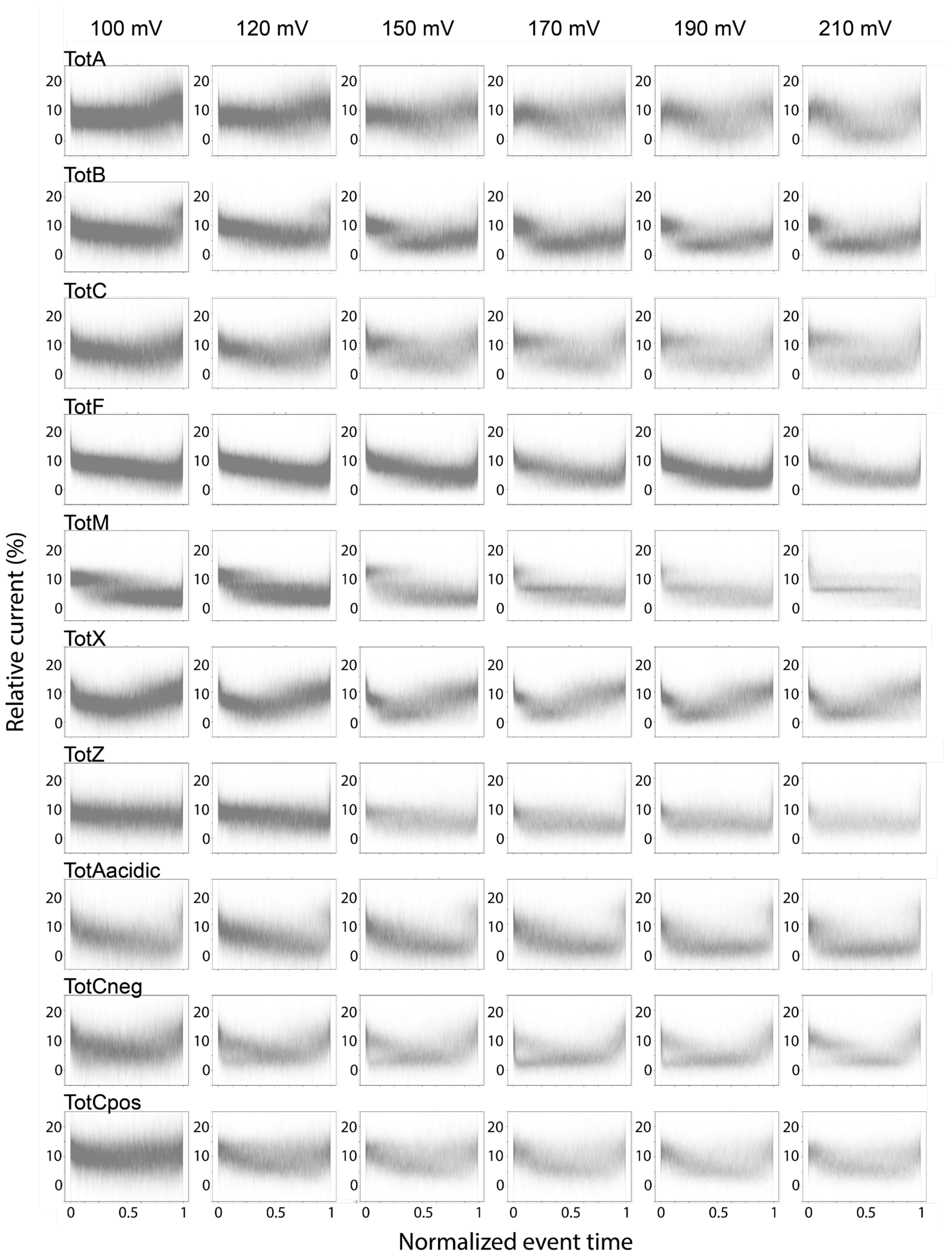


Figure S12. Time normalized overlay plots of randomly chosen events recorded various voltages for each Tot family protein.

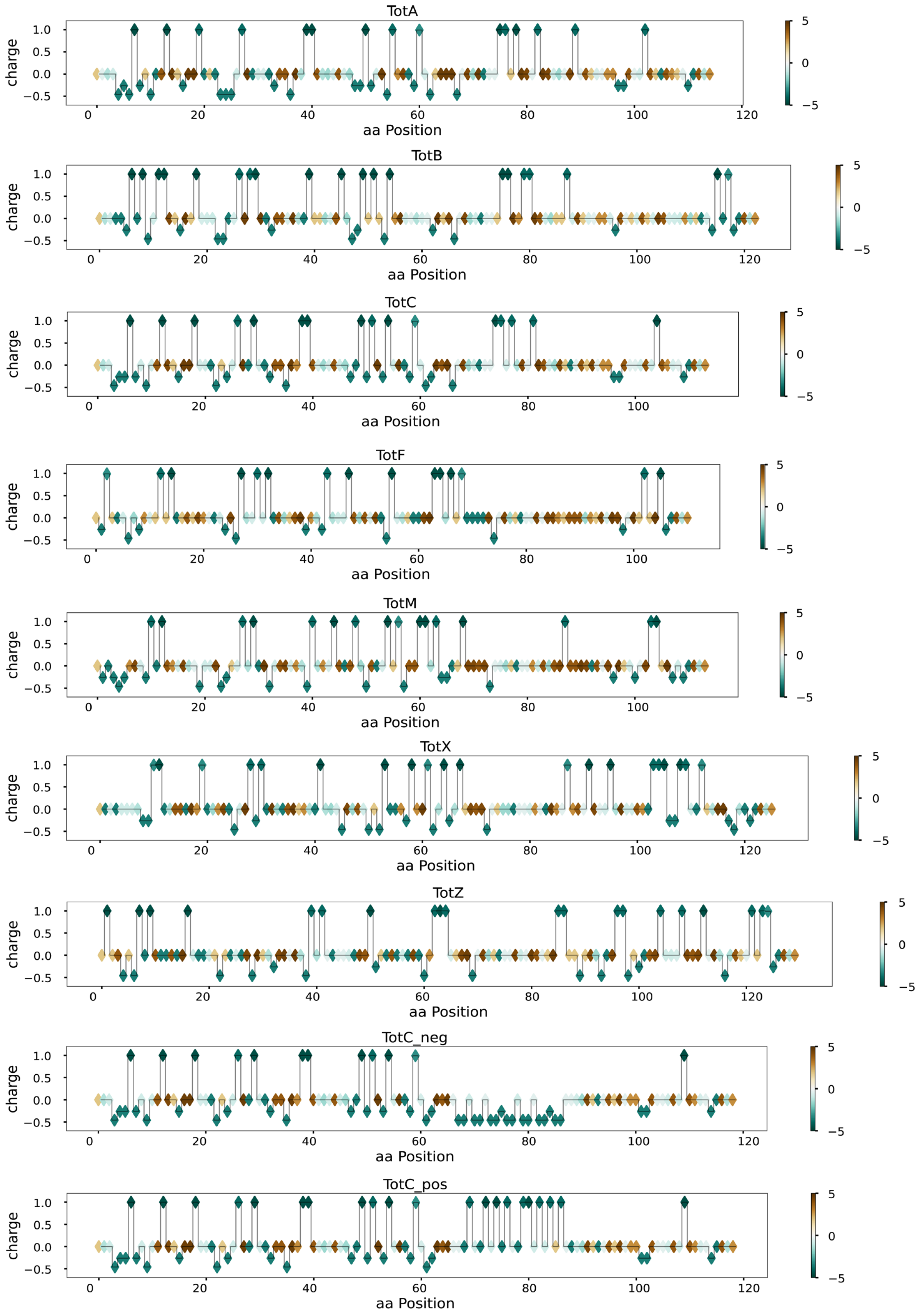


Figure S13. Charge and hydropathy properties along the aa residues of the analyzed proteins.

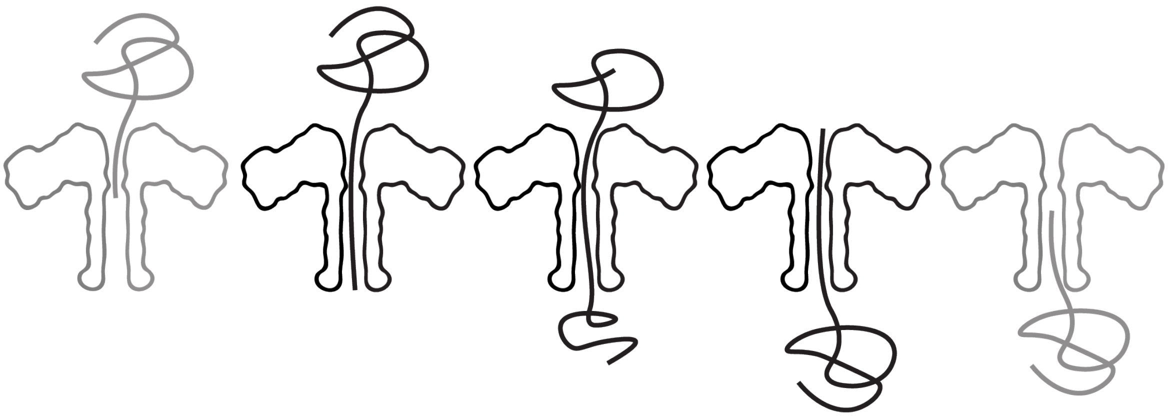

Figure S14. Illustration of the protein translocation scenario in aerolysin nanopore. In the sliding window approach, the beginning and end (gray) of the process are not reflected.

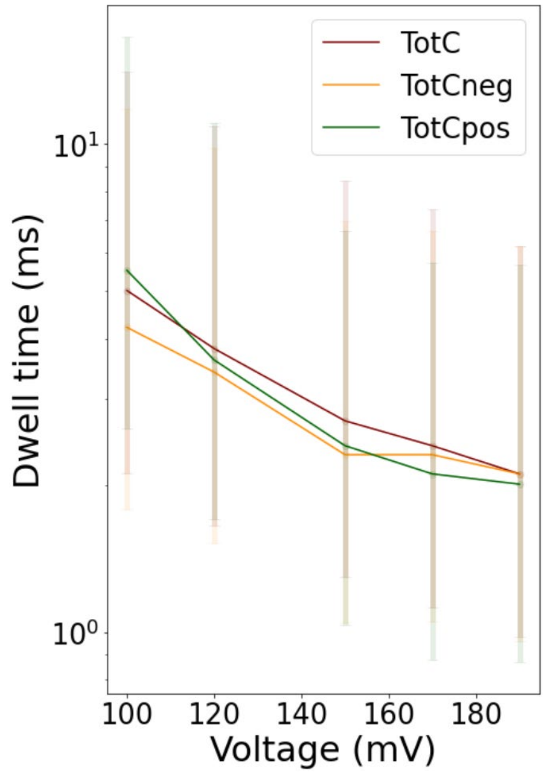


Figure S15. Population dwell times of each protein were fitted with the Fokker Planck equation and plotted against applied potential. Maximum and halfwidth of the distributions are shown.
